# Two marine GH29 *α*-L-fucosidases from an uncultured *Paraglaciecola* sp. specifically hydrolyze fucosyl-*N*-acetylglucosamine regioisomers

**DOI:** 10.1101/2021.06.01.446583

**Authors:** Mikkel Schultz-Johansen, Peter Stougaard, Birte Svensson, David Teze

## Abstract

l-Fucose is the most widely distributed l-hexose in marine and terrestrial environments, and presents a variety of functional roles. l-Fucose is the major monosaccharide in the polysaccharide fucoidan from cell walls of brown algae, and is found in human milk oligosaccharides and the Lewis blood group system, where it is important in cell signaling and immune response stimulation. Removal of fucose from these biomolecules is catalyzed by fucosidases belonging to different carbohydrate-active enzyme (CAZy) families. Fucosidases of glycoside hydrolase family 29 (GH29) release *α*-l-fucose from non-reducing ends of glycans and display activities targeting different substrate compositions and linkage types. While several GH29 fucosidases from terrestrial environments have been characterized, much less is known about marine members of GH29 and their substrate specificities, as only four marine GH29 enzymes were previously characterized. Here, five GH29 fucosidases originating from an uncultured fucoidan-degrading marine bacterium (*Paraglaciecola* sp.) were cloned and produced recombinantly in *E. coli.* All five enzymes (Fp231, Fp239, Fp240, Fp251, Fp284) hydrolyzed the synthetic substrate CNP-*α*-l-fucose. By screening each of these enzymes against up to 17 fucose-containing oligosaccharides Fp231 and Fp284 showed strict substrate specificities against the fucosyl-*N*-acetylglucosamine regioisomers Fuc(*α*1,4)GlcNAc and Fuc(*α*1,6)GlcNAc, respectively, the former representing a new specificity. Fp231 is a monomeric enzyme with pH and temperature optima at pH 5.6–6.0 and 25°C, hydrolyzing Fuc(*α*1,4)GlcNAc with *k*_cat_ = 1.3 s^−1^ and *K*_m_ = 660 μM. Altogether, the findings extend our knowledge about GH29 family members from the marine environment, which are so far largely unexplored.

## INTRODUCTION

Fucose in several ways is an unusual sugar; the l-configuration, the deoxy group (6-deoxy-galactose), and the exclusive participation in *α*-glycosidic bonds despite having an equatorial 2-OH group. Yet, fucose is the most common deoxy sugar, and the most common l-sugar in the biosphere. As a 6-deoxy sugar, it lacks the primary hydroxyl group that holds a high degree of freedom, and is more hydrophobic than most other monosaccharides. This is an important feature in protein-carbohydrate recognition, and hence fucose is found in a wide range of organisms (Staudacher et al. 1999). In humans, fucose residues are usually unsubstituted, both on *N*- and *O*-glycans, and on the Lewis blood group antigens, which are cell recognition sites for numerous viral and bacterial pathogens (Boren et al. 1993; Kubota et al. 2012; Schneider et al. 2017). Notably, Lewis antigen oligosaccharide structures are found in mucin – the protective mucus barrier covering the gastrointestinal tract (Tailford et al. 2015) – and on human milk oligosaccharides (HMOs) (Ayechu-Muruzabal et al. 2018). Some gut microbes are able to degrade and utilize mucin and HMOs for growth (Sela et al. 2008; Marcobal et al. 2011), which can have positive and negative consequences to the host. In bacteria, fucose is a prominent sugar in oligosaccharide motifs of surface glycans (Mäki and Renkonen 2003). Moreover, fucose plays a structural role in fucoidans, which are highly sulfated and substituted fucose polymers found in cell walls of brown algae and in some marine invertebrates (Deniaud-Bouet et al. 2017).

Glycoside hydrolases (GHs) acting on fucosidic bonds belong to glycoside hydrolase families GH29, GH95, GH107, GH139, GH141, GH151 or GH168 in the CAZy database (www.cazy.org; (Lombard et al. 2014)). While enzymes of GH107 and GH168 are *endo-*fucosidases hydrolyzing fucoidan, a fucan present in brown algae (Colin et al. 2006; Schultz-Johansen et al. 2018; Vickers et al. 2018) and marine invertebrates (Shen et al. 2020), respectively, the other families comprise *exo*-fucosidases. GH95 mainly contains *α*-1,2-l-fucosidases, one characterized member in each of GH139 and GH141 is targeting specific *α*-l-fucose motifs in pectin (Ndeh et al. 2017), and enzymes from GH151 are not yet characterized in detail (Sela et al. 2012; Lezyk et al. 2016). By contrast, GH29, the largest family of fucosidases, covers a broad spectrum of substrate specificities. Some GH29 fucosidases act on fucosyl-*N*-acetylglucosamine (Fuc(*α*1,3/4/6)GlcNAc) motifs found in *N*-glycans, Lewis blood group antigens and HMOs. Some GH29 enzymes are highly specific for one linkage type, as for example AlfB and AlfC from *Lactobacillus casei* hydrolyzing Fuc(*α*1,3)GlcNAc and Fuc(*α*1,6)GlcNAc, respectively (Rodriguez-Diaz et al. 2011), while others are able to react on several linkage types (Ashida et al. 2009; Hobbs et al. 2019).

Most characterized GH29 fucosidases are of terrestrial origin while only a few are from marine organisms (Tarling et al. 2003; Dong et al. 2017; Ono et al. 2019; Hong et al. 2021). Terrestrial and marine glycans have distinctly different compositions, and especially algal cell walls contain polysaccharides (e.g. fucoidan) not found in organisms living on land (Popper et al. 2011). Indeed, the function of fucosidases in the marine environment is less described, although enzymes of GH29 are key in degradation of fucoidan, as exemplified by the marine bacterium *Lentimonas* sp. CC4. This specialized fucoidan-degrader encodes 35 phylogenetically diverse GH29 enzymes likely encompassing different substrate specificities connected with the complexity of glycan motifs in fucoidan structures (Sichert et al. 2020). Investigations of marine GH29 enzymes, offer an opportunity for identification of new substrate specificities not covered by currently characterized GH29 enzymes.

Previously, metagenome data revealed seven putative GH29 fucosidases encoded by a fucoidan-degrading uncultured marine bacterium (*Paraglaciecola* sp.) alongside with three GH107 *endo*-1,4-fucanases acting on fucoidan (Schultz-Johansen et al. 2018). Here, the five GH29 enzymes, Fp231, Fp239, Fp240, Fp251, Fp284, which are encoded closest in the genome to the *endo*-1,4-fucanases were cloned, produced recombinantly in *E. coli* and analyzed for *exo*-*α*-l-fucosidase activity on a comprehensive collection of oligosaccharides. Three of these enzymes, Fp239, Fp240, Fp251, were only active on the artificial substrate 2-chloro-4-nitrophenyl-*α*-l-fucopyranoside (CNP-Fuc), whereas the disaccharide Fuc(*α*1,6)GlcNAc was selectively hydrolyzed by Fp284, similarly to the activity of AlfC from *L. casei* (Rodriguez-Diaz et al. 2011; Klontz et al. 2020), and Fuc(*α*1,4)GlcNAc by Fp231. Fp231 represents a new substrate specificity of strict regioselectivity for 1,4-linked fucosyl-*N*-acetylglucosamine, and the enzyme was further characterized biochemically. Altogether the findings provide new information about the GH29 family and highlight the marine environment as a prospective source of undiscovered enzyme activities.

## RESULTS

### Heterologous production and substrate screening of five marine GH29 enzymes

Five putative GH29 fucosidases were identified in a partial genome sequence of a fucoidan-degrading bacterium previously obtained by using a metagenomics strategy. The genes encoding these GH29s (Fp231, Fp239, Fp240, Fp251 and Fp284) were flanked by GH107 fucoidanases and other *α*-l-fucosidases belonging to GH95 and GH141 (Schultz-Johansen et al. 2018). According to protein domain structure predictions, the five GH29 enzymes all carry a secretion signal peptide (**Figure S1**). Fp231, Fp240 and Fp251 contain a C-terminal domain, believed to be involved in carbohydrate-binding in GH29 fucosidases (Klontz et al. 2020), whereas Fp239 and Fp284 are shorter single-domain enzymes. The full-length sequence of each of the five enzymes without predicted signal peptides was cloned and expressed in *E. coli* BL21(DE3) yielding soluble protein in cell lysates for Fp239, Fp240, Fp251 and Fp284 purified using IMAC (**Figure S2**). However, protein corresponding to Fp231 was not detected neither in the soluble nor in the insoluble fractions of cell lysates. Attempted expression of shorter versions of the Fp231 gene (**Supplementary Table I**) gave a similar dissatisfying outcome. Perhaps, as growth and growth rate of the induced host cells decreased, Fp231 is toxic to *E. coli*. However, expressing the full-length gene including the predicted N-terminal signal peptide produced soluble Fp231 detected in the culture supernatant, indicating that the native secretion signal functioned in *E. coli* allowing secretion of Fp231.

The *exo*-*α*-l-fucosidase activity of Fp231, Fp239, Fp240, Fp251 and Fp284 was confirmed using the substrate CNP-Fuc. Since these GH29 enzymes originated from a fucoidan-degrader they may be involved in the final degradation steps of this fucose-containing polysaccharide. Unfortunately, well-defined oligosaccharides derived from fucoidan are not available from commercial suppliers and would have to be prepared either by chemical means (Khatuntseva et al. 2000; Krylov et al. 2011) or by partial enzymatic digestion of the natural polysaccharides (Dong et al. 2017; Silchenko et al. 2017). Therefore, the five GH29 enzymes were screened against a collection of 17 fucose-containing oligosaccharides, found in HMOs and blood group antigens and representing a variety of substrate lengths, configuration and linkage types (**Figure 1**). For Fp239, Fp240 and Fp251, no activity was detected on any of these oligosaccharides, and CNP-Fuc was the only hydrolyzed substrate. By contrast, Fp231 and Fp284 hydrolyzed fucosyl-*N*-acetylglucosamine disaccharides, albeit with very narrow substrate specificity, thus of Fp231 for Fuc(*α*1,4)GlcNAc and of Fp284 for Fuc(*α*1,6)GlcNAc (**Figure 1**). Specificity against Fuc(*α*1,6)GlcNAc was previously reported for the GH29 fucosidase AlfC from *L. casei* (Rodriguez-Diaz et al., 2011), and the specificity of Fp284 is similar to that of AlfC, which also has no or negligible activity on fucosyl-*N*-acetylglucosamine disaccharides with other linkage types. Conversely, the strict substrate specificity of Fp231 against Fuc(*α*1,4)GlcNAc is yet unreported, hence Fp231 was subjected to more detailed biochemical characterization.

**Figure 1:**
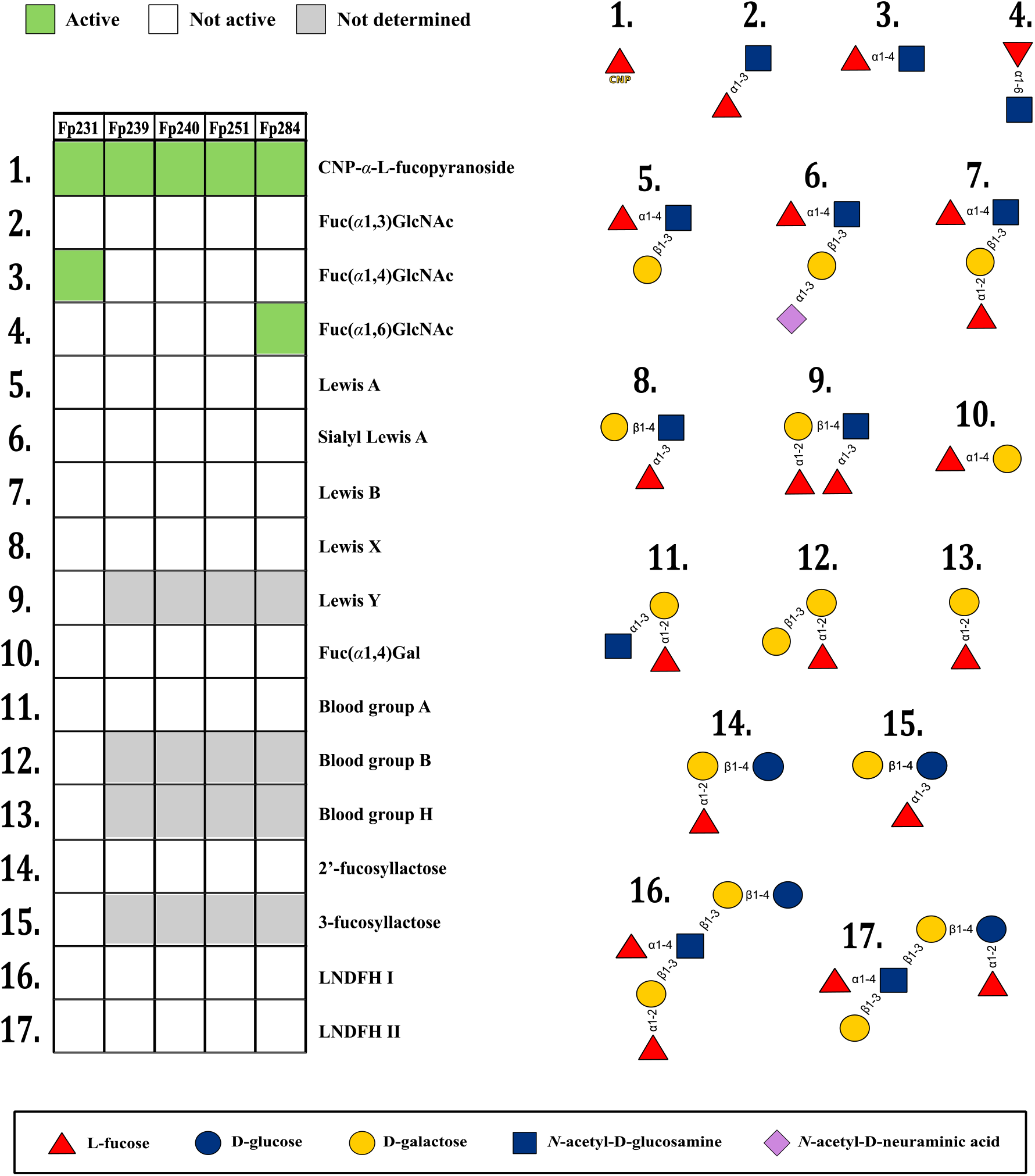
Substrate screening of GH29 fucosidases identified from *Paraglaciecola* sp. in a marine metagenome. The five enzymes were screened against up to 17 oligosaccharides in a substrate collection and activity was assessed by HPAEC-PAD, fluorescence-assisted carbohydrate electrophoresis or absorbance measurements at 410 nm. Structural representation of oligosaccharides follows the SNFG (Symbol Nomenclature for Glycans) system (Varki et al. 2015).

### Biochemical characterization of GH29 Fp231

Recombinant Fp231 was produced in *E. coli* as described above, concentrated in the cell free culture supernatant and purified to homogeneity by IMAC (yield 2 mg·L^-1^ culture) followed by SEC (**Figure 2A**). An estimated molecular mass of 52 kDa was found from SEC (**Figure 2B**), hence Fp231 is a monomer in solution (theoretical molecular mass is 56.2 kDa).

**Figure 2:**
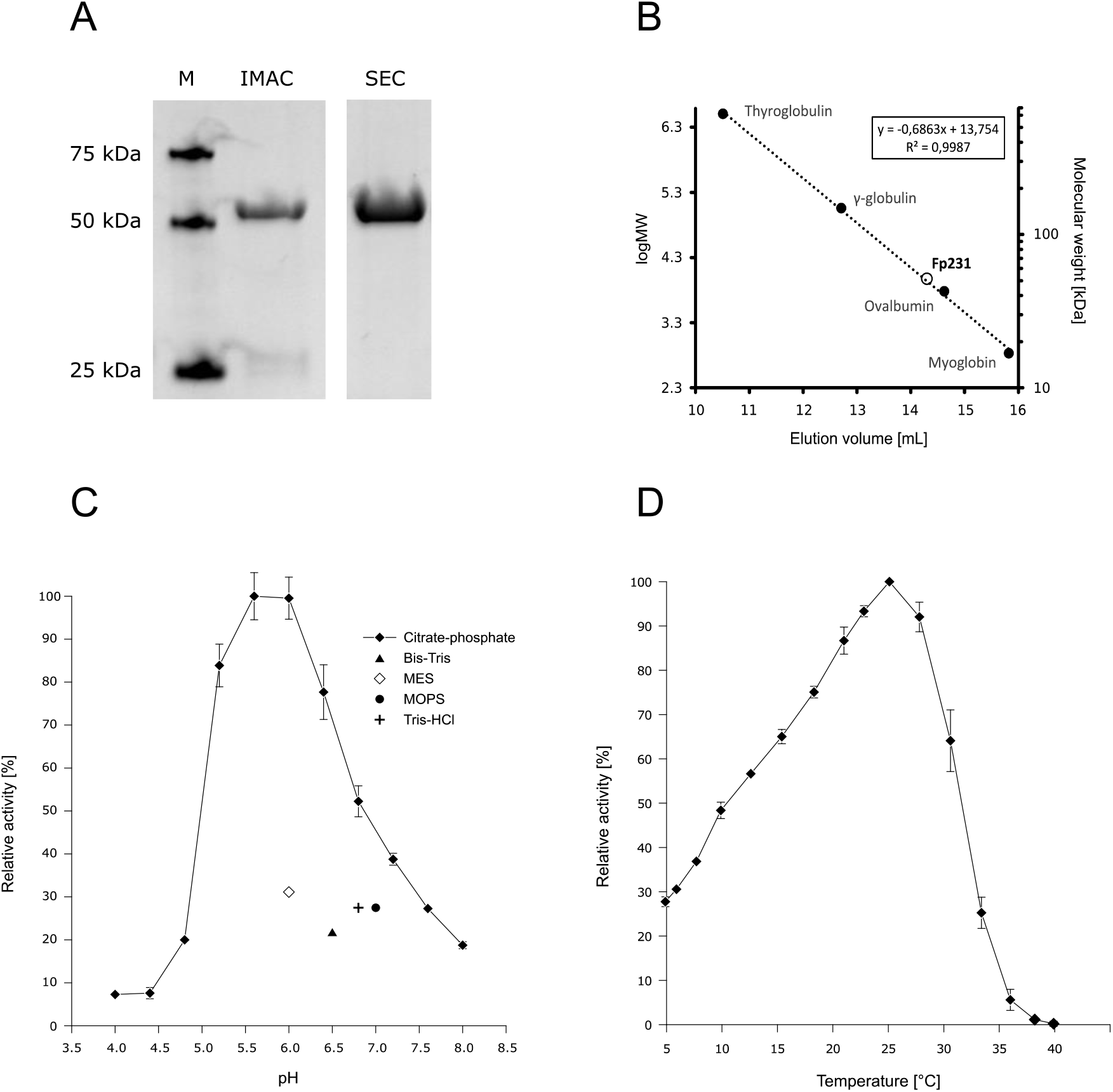
Purification and biochemical characterization of recombinant Fp231. (**A**) SDS-PAGE of Fp231 purified by immobilized metal affinity chromatography (IMAC) and size exclusion chromatography (SEC; Sephacryl S-200 HR), and compared to a protein marker (M). (**B**) Analytical size exclusion chromatography (ENrich™ SEC 650 column) of purified Fp231. The molecular mass was estimated from a calibration curve produced in a separate run with the markers thyroglobulin (670 kDa), *γ*-globulin (158 kDa), ovalbumin (44 kDa), and myoglobin (17 kDa). (**C**) Optimum pH of Fp231 was determined at room temperature in reactions containing 1 mM 2-chloro-4-nitrophenyl-*α*-l-fucopyranoside in 50 mM sodium citrate-phosphate (pH 4.0 – 8.0); 50 mM Bis-Tris (pH 6.5); 50 mM MES (pH 6.0); 50 mM MOPS (pH 7.0) and 50 mM Tris-HCl (pH 6.8). (**D**) Optimum temperature of Fp231 was determined towards 1 mM 2-chloro-4-nitrophenyl-*α*-l-fucopyranoside in 50 mM sodium citrate-phosphate pH 6.0. Maximum activity is set at 100%; error bars represent the standard deviation of triplicate assays.

CNP-Fuc a chlorinated version of the synthetic substrate 4-nitrophenyl-*α*-l-fucopyranoside (*p*NP-Fuc) allows continuous monitoring of hydrolysis at slightly acidic pH, as p*K*_a_ of CNP is 5.43 compared to *p*NP having p*K*_a_ = 7.24. CNP-Fuc was used to determine pH and temperature optima, as well as effects of NaCl and metal ions on the activity. The activity of Fp231 was optimal at pH 5.6 – 6.0 in 50 mM citrate-phosphate; in other buffers the activity was reduced (**Figure 2C**). Importantly, Fp231 has cold-active characteristics (Gerday et al. 1997) and shows temperature optimum for activity at about 25°C, retains about 50 % of this activity at 10°C, and is quickly inactivated over 30°C (**Figure 2D**). Yet, purified Fp231 remains active after storage for >1 month at 4°C in 50 mM citrate-phosphate pH 6.0. The effect of metal ions on Fp231 was tested in 50 mM Bis-Tris pH 6.5, to avoid chelating citrate and phosphate ions, indicating sensitivity to Co^2+^ and Ni^2+^ and almost complete inhibition by Zn^2+^. Notably, Fp231 was activated by 1 mM EDTA, but no effect was observed at 5 mM EDTA (**Table I**) and there was little effect of 0 – 2 M NaCl on activity, where Fp231 maintained > 70 % activity, with maximum at 0.2 – 0.3 M NaCl (**Figure S3**).

**Table I:** Effect of metal ions and EDTA on the activity of Fp231 toward CNP-Fuc. Values are the mean of triplicates with S.D. <1.5%.

| Effector | Relative activity (%) |  |
| --- | --- | --- |
|  | 1 mM | 5 mM |
| Ca<sup>2+</sup> | 105 | 104 |
| Mg<sup>2+</sup> | 89 | 100 |
| Zn<sup>2+</sup> | 9 | 1 |
| Co<sup>2+</sup> | 52 | 46 |
| Ni<sup>2+</sup> | 41 | 23 |
| Mn<sup>2+</sup> | 82 | 81 |
| Li<sup>+</sup> | 103 | 98 |
| K<sup>+</sup> | 99 | 95 |
| EDTA | 166 | 102 |

In the initial substrate screening, activity against Fuc(*α*1,4)GlcNAc was monitored by high pressure anion exchange chromatography coupled with pulsed amperometric detection (HPAEC-PAD), showing hydrolysis of Fuc(*α*1,4)GlcNAc to fucose and GlcNAc after 6 h at 37°C (**Figure 3A**). To verify this specificity, Fp231 was mixed with oligosaccharide substrate candidates and incubated 24 h at the optimal temperature (25°C) and the reaction products were analyzed by fluorophore-assisted carbohydrate electrophoresis (FACE) compared with oligosaccharides without addition of Fp231 (**Figure 3B**). Under these conditions, Fp231 completely hydrolyzed Fuc(*α*1,4)GlcNAc, whereas Fuc(*α*1,3)GlcNAc and Le^a^ trisaccharide (Gal(β1,3)[Fuc*α*1,4]GlcNAc) were not degraded. The kinetic constants for hydrolysis of CNP-Fuc and Fuc(*α*1,4)GlcNAc were determined by Michaelis-Menten analyses (**Table II**, **Figure S4**). *K*_m_ values of Fp231 for CNP-Fuc and Fuc(*α*1,4)GlcNAc are in the usual range for GH29 fucosidases. According to the BRENDA database (as of March 23^rd^, 2021), 53 out of 70 characterized GH29 fucosidases display submillimolar *K*_m_ against *p*NP-*α*-l-Fuc, www.brenda-enzymes.org, (Chang et al. 2021). The relatively larger *k*_cat_ observed for Fp231 towards CNP-Fuc is also common for substrates having leaving groups with low p*K*_a_ (7.24 and 5.43 for *p*NP and CNP, respectively, compared to ≈16 for an equatorial OH group on a sugar ring).

**Figure 3:**
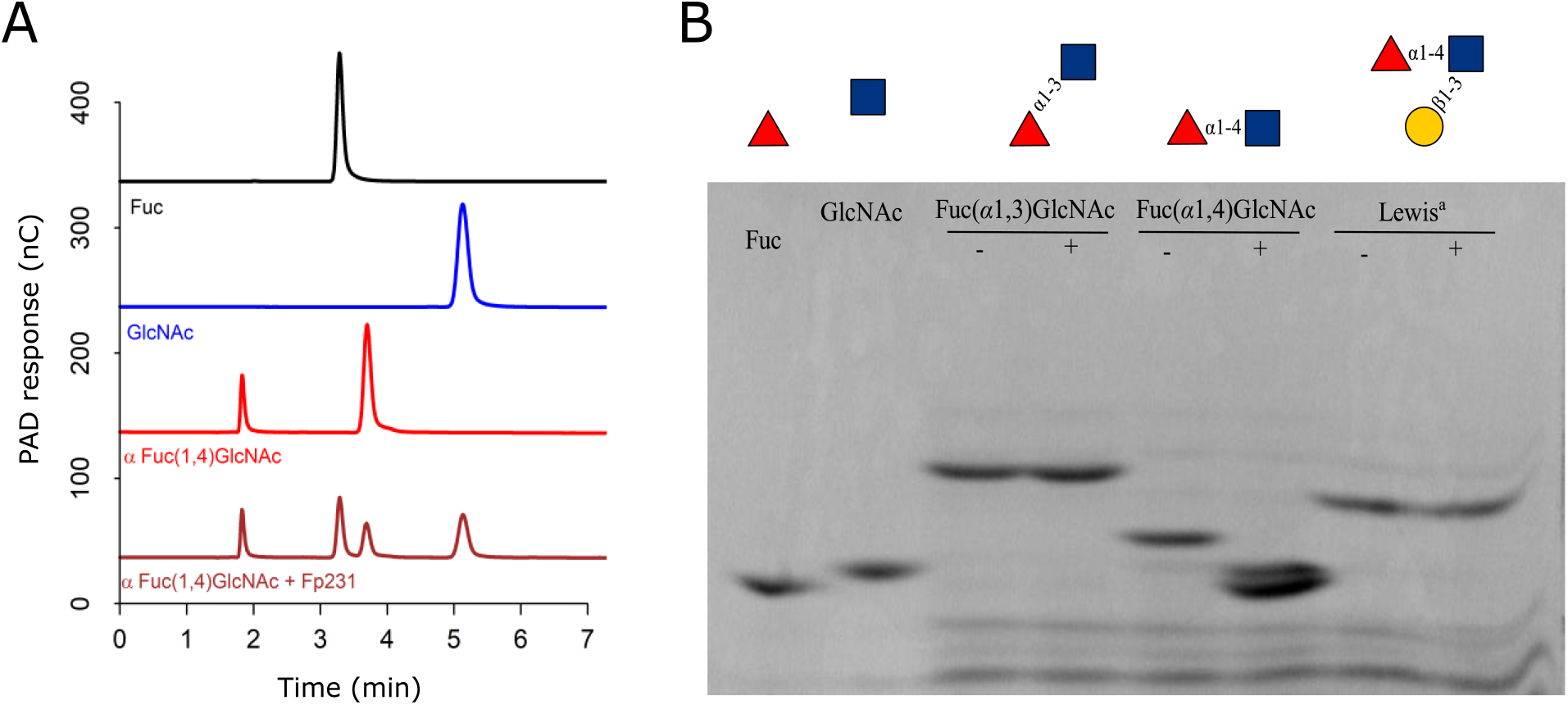
Hydrolytic activity of Fp231 on oligosaccharides. (**A**) HPAEC-PAD chromatograms showing the hydrolysis of 2 mM Fuc(*α*1,4)GlcNAc by 0.2 μM Fp231 for 6 h at 37°C. (**B**) Fluorescence-assisted carbohydrate electrophoresis (FACE) gels products formed by 5 μM Fp231 from 10 μg oligosaccharide (100 μL, 25°C, ON). Incubation without (-) Fp231 served as control. The reaction products were fluorophore labeled and separated in acrylamide gels. Enzyme activity was evaluated as a change in mobility of the fluorescent oligosaccharides. Monosaccharide standards (Fuc and GlcNAc) were included for comparison.

**Table II:** Kinetic constants obtained for Fp231

| Substrate | $K_m$ (mM) | $k_{cat}$ (s<sup>-1</sup>) | $k_{cat}/K_m$ (s<sup>-1</sup> mM<sup>-1</sup>) |
| --- | --- | --- | --- |
| CNP-Fuc | 0.14 ± 0.01 | 31 ± 0.5 | 221 |
| Fuc(α1,4)GlcNAc | 0.66 ± 0.05 | 1.3 ± 0.04 | 1.97 |

Members of GH29 are retaining enzymes and some are able to catalyze transglycosylation leading to the formation of new glycosidic bonds (Rodriguez-Diaz et al. 2013; Lezyk et al. 2016; Zeuner et al. 2019). Transglycosylation can thus occur in reactions with a wild type glycoside hydrolase, or be enhanced through e.g. mutagenesis, change of reaction conditions, increasing the substrate concentration and so forth (Bissaro et al. 2015; Zeuner et al. 2019; Teze et al. 2021). In this context, AlfB and AlfC from *L. casei* by reaction with *p*NP-*α*-l-fucose and GlcNAc generated Fuc(*α*1,3)GlcNAc and Fuc(*α*1,6)GlcNAc, respectively, which are important building blocks in HMOs or found in glucoconjugates at mucosal surfaces (Rodriguez-Diaz et al. 2013; Teze et al. 2021). Fuc(*α*1,4)GlcNAc has similar biological importance, and therefore Fp231 was investigated for ability to catalyze formation of Fuc(*α*1,4)GlcNAc and Fuc(*α*1,4)GlcNAc-linked products. Unfortunately, no transglycosylation products were observed when using wild type Fp231 with 20 mM CNP-Fuc as donor and 200 mM GlcNAc as acceptor (data not shown). In an attempt to stimulate transglycosylation, Fp231 mutants H174F and W225H were designed by using mutant transfer based on a strategy of conserved-residue mutation, which enabled in this case increased transglycosylation by both *L. casei* AlfB and AlfC of GH29 (Teze et al. 2021). However, still no transglycosylation product was obtained.

### Sequence analysis

The sequence of Fp231 was compared to sequences available in public databases using BLASTp searches. The identified sequence homologs were derived from marine bacteria, uncharacterized proteins from *Paraglaciecola* sp. L3A3 (WP 158972757.1) and *Thalassotalea* sp. HSM 43 (WP 135437836.1) being the two closest relatives of 75 % and 71 % identity to Fp231, respectively. The sequence of Fp231, Fp239, Fp240, Fp251 and Fp284 were compared with sequences of characterized GH29 enzymes (42 GH29 members characterized as of May 27^th^, 2021) by multiple sequence alignment and phylogenetic analysis (**Figure 4**). Notably, enzymes of family GH29 occur in two phylogenetic clades reflecting their substrate specificity (Sakurama et al. 2012). Members of clade A hydrolyze *p*NP-Fuc (CNP-Fuc is used in the present study) and have a relatively narrow substrate range, while members of clade B often display broader substrate specificity, but are inactive on *p*NP-Fuc. The five GH29 described here all cluster with clade A enzymes, in accordance with their activity on CNP-Fuc and narrow substrate specificity as verified by lack of action on a series of oligosaccharides (**Figure 1**).

**Figure 4:**
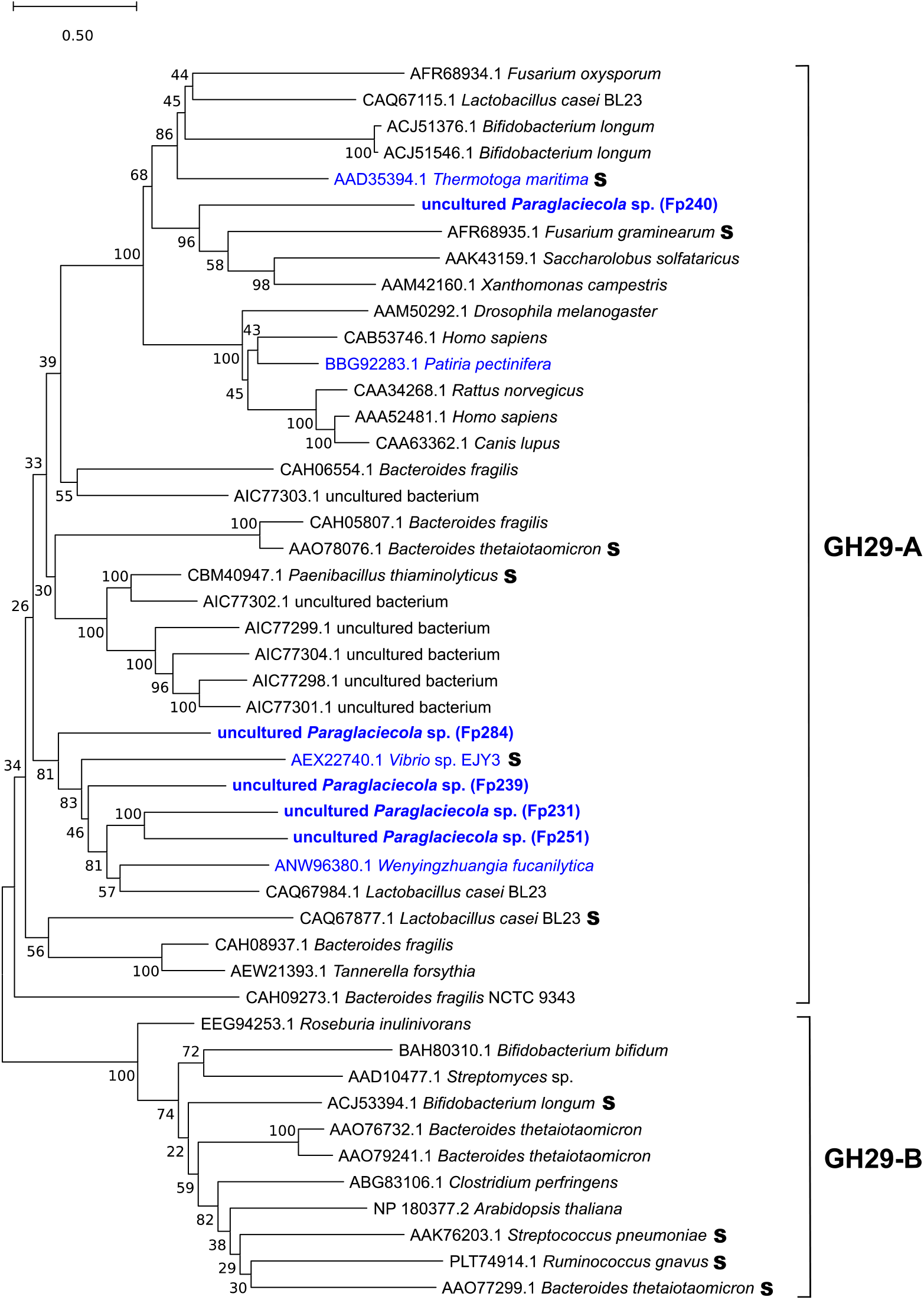
Phylogenetic relationship of characterized GH29 fucosidases. Neighbor-joining phylogenetic tree showing the relationship between the five GH29s from the present study (bold font) and previously characterized GH29 fucosidases from the CAZy database. Accession number and organism are listed for each protein. **S** indicates characterized GH29 members that have been structure-determined and proteins of a marine origin is indicated in blue. Proteins belonging to clade GH29-A and GH29-B are indicated (Sakurama et al., 2012). The scale bar represents the number of substitutions per site.

Four out of the five GH29 enzymes in the present study, Fp231, Fp239, Fp251 and Fp284, cluster in the phylogenetic tree together with two other characterized enzymes Alf1_Wf from the marine bacterium *Wenyingzhuangia fucanilytica*, and AlfC from the terrestrial *L. casei.* Notably, both Alf1_Wf and AlfC are active on fucose linked to GlcNAc. Dong and coworkers (2017) reported Alf1_Wf to release fucose very slowly from the Lewis^a^ trisaccharide (Gal β1,3[Fuc*α*1,4]GlcNAc), while AlfC has specificity for Fuc(*α*1,6)GlcNAc (Rodriguez-Diaz et al., 2011). Interestingly, Alf1_Wf is able to release fucose from oligosaccharides obtained by *endo*-fucoidanase degradation of fucoidan (Dong et al. 2017). Since Fp231, Fp239, Fp251 and Fp284 stem from a fucoidan-degrading bacterium and cluster with Alf1_Wf in the phylogenetic tree, it is possible that they share functional role and mechanism. However, Fp231 was unable to release fucose from HMW fucoidan (>10 kDa) from the macroalgae *Fucus vesiculosus, Fucus serratus* and *Ascophyllum nodosum* or from LMW fucoidan (<10 kDa) from *F. vesiculosus* (**Figure S5**). Fp240 groups with other characterized GH29 members with the nearest homolog being from *F. graminearum*.

To determine the structure of Fp231, crystallization was attempted in 96-well plate format using 1248 conditions in commercial factorial screen kits. However, no crystal formation was observed.

## DISCUSSION

Lately, attention has been drawn to the GH29 family and the collection of active enzymes has been expanded (Grootaert et al. 2020). Yet, little is known about marine GH29 enzymes, and here we show that new specificities are found in the marine environment. Specifically, the enzyme Fp231 presents a new substrate specificity as it degrades Fuc(*α*1,4)GlcNAc, while remaining inactive on other fucosyl-*N*-acetylglucosamine disaccharides and on longer oligosaccharides including such containing the Fuc(*α*1,4)GlcNAc motif. Conversely, the already characterized GH29 enzymes AlfB and AlfC have specificity for two other Fuc-GlcNAc disaccharides, Fuc(*α*1,3)GlcNAc and Fuc(*α*1,6)GlcNAc (Rodriguez-Diaz et al. 2011), respectively. Thus, with the Fp231 described here included, all possible strict specificities against reducing Fuc-GlcNAc disaccharides are known.

Strictly regiospecific enzymes are attractive for HMO synthesis (Zeuner et al. 2019), but application of Fp231 in such processes seems questionable, as it did not catalyze formation of transglycosylation products. Moreover, we so far have not obtained crystals of Fp231, making it impossible to elucidate molecular determinants of the Fuc(*α*1,4)GlcNAc specificity. It has been proposed that the specificity of AlfC Fuc(*α*1,6)GlcNAc stems from an aromatic subsite +1 accommodating the GlcNAc moiety, in combination with a dynamic loop that opens and closes the active site (Klontz et al. 2020). As AlfC, the closest structure determined homolog of Fp231 (42% sequence identity), has a reported flexible active site within a functional tetramer, while Fp231 is monomeric, homology modelling cannot reliably identify specificity determinants.

A limitation of the present study is that marine oligosaccharides are essentially not available and the enzymes therefore were screened for activity against oligosaccharides associated with terrestrial sources. Fp231 released no fucose when incubated with native marine fucoidan polysaccharides (**Figure S5**). Fucoidans are structurally highly diverse glycans, which are sulfated and acetylated, and apart from fucose contain other monosaccharides e.g. galactose, mannose, xylose, galacturonic acid, glucuronic acid and glucosamine, albeit in smaller amounts (Sichert et al. 2021). Therefore, it is likely that the action of additional depolymerizing or debranching enzymes is required for Fp231 to access substrate bonds. It was demonstrated that depolymerization of fucoidan by specific *endo*-hydrolases can be required to detect GH29 activity (Dong et al. 2017). Moreover, the fucoidan-degrading enzymatic cascade in marine bacteria is complex and can involve hundreds of CAZymes including fucosidases, sulfatases and deacetylases (Sichert et al. 2020). The fact that such supplementing enzyme activities are encoded in the same genomic region as Fp231 (Schultz-Johansen et al. 2018), suggests that Fp231 may degrade oligosaccharides released following a defined enzymatic cascade.

*α*-fucans represent a major glycan motif among fucoidans (Deniaud-Bouet et al. 2017), and therefore Fuc(*α*1,4)Fuc in particular would have been a logic choice to include in the substrate screening of Fp231, had it been available. On the other hand, as Fuc(*α*1,4)-linked galactose was not degraded by Fp231, it seems unlikely that Fp231 would be active on Fuc(*α*1,4)Fuc due to the similar di-axial configuration of these two disaccharides, compared to the axial-equatorial bond cleaved by Fp231 (**Figure S6**). As such, Fp231 appear to target Fuc(*α*1,4)GlcNAc motifs.

According to the Carbohydrate Structure Database (CSDB), the Fuc(*α*1,4)GlcNAc disaccharide (CSDB code aLFuc(1-4)[Ac(1-2)]bDGlcN) is part of 55 glycan motifs (Toukach and Egorova 2015). Remarkably, it appears to cover a range of functions in vastly different organisms, such as *O*-antigens of bacteria e.g. *Escherichia, Shigella, Helicobacter* or *Salmonella* genera, in plant glycoproteins (Melo et al. 1997; Fitchette et al. 1999; Strasser et al. 2007) and in HMOs or Lewis^a^ antigens in humans (Green 1989; Ayechu-Muruzabal et al. 2018). Given the variety of terrestrial biotopes, the present finding of a strictly specific marine GH29 enzyme, makes us speculate if the Fuc(*α*1,4)GlcNAc motif is also prevalent in marine environments. Indeed, both fucose and *N*-acetylglucosamine are abundant in the ocean (as fucoidan and chitin), however, observations of this motif is yet unreported.

## MATERIALS & METHODS

### Chemicals

Fuc(*α*1,3)GlcNAc (A151500), Fuc(*α*1,4)GlcNAc (A151550), Fuc(*α*1,6)GlcNAc (A151570), Fuc(*α*1,4)Gal (F823520), Lewis^a^ trisaccharide (L394000) and Sialyl Lewis^a^ (S397500 were purchased from Toronto Research Chemicals (North York, ON, Canada); 2-chloro-4-nitrophenyl-*α*-L-fucopyranoside (EC05990), Lewis^b^ (OL02434), Lewis^x^ (OL06490), Lewis^y^ (OL06521), blood group A trisaccharide (OB04438), blood group B trisaccharide (OB01663), blood group H disaccharide (OB05907), 2’-fucosyllactose (OF06739), LNDFH I (OL01664) and LNDFH II (OL06826) from Carbosynth Ltd (Compton, Berkshire, UK); and 3-fucosyllactose (GLY060) from Elicityl (Crolles, France). Fucoidans from *Fucus serratus* (YF09360) and *Ascophyllum nodosum* (YF09363) were from Carbosynth and fucoidan from *Fucus vesiculosus* (F8190) was from Sigma. HMW and LMW *F. vesiculosus* fucoidan were separated by passing an aqueous fucoidan solution over a 10 kDa cut-off ultrafiltration membrane (Merck) followed by lyophilization.

### Sequence analysis

Metagenome sequence data (Schultz-Johansen et al. 2018) were downloaded from the MG-RAST server: https://www.mg-rast.org (study no. mgp84310). Nucleotide sequences of Fp231, Fp239, Fp240, Fp251 and Fp284 were retrieved from the assembled metagenome and deposited in GenBank under accessions MW623630, MW623631, MW623632, MW623633, and MW623634. Protein sequences of the five GH29 encoded enzymes and those of characterized GH29 members were compared (http://www.cazy.org/, as of 27^th^ May 2021) by alignment using CLUSTAL Omega (Madeira et al. 2019) and constructing the corresponding phylogenetic trees using the neighbor-joining algorithm and Jones-Taylor-Thornton model in MEGA X (Kumar et al. 2018). The final tree was based on 100 bootstrap replicates.

Signal peptide prediction was performed using SignalP 5.0 (Almagro Armenteros et al. 2019). The molecular mass and extinction coefficient of Fp231 (without signal peptide) were calculated to 56.2 kDa and 110935 M^-1^·m^-1^ respectively using ProtParam: https://web.expasy.org/protparam/

### Cloning

Novagen® expression vectors pET9a and pET15b, carrying kanamycin and ampicillin resistance genes respectively, were obtained from Merck. Genes encoding Fp231, Fp239, Fp240, Fp251 and Fp284 were amplified by PCR from metagenomic DNA isolated previously (Schultz-Johansen et al. 2018). Primers were constructed either with or without the predicted secretion signal sequences and primer overhangs were added to allow fusion with pET9a and pET15b vector fragments based on the USER™ fusion cloning strategy as described by Geu-Flores et al, 2007 (**Supplementary Table I**). Assembled GH29 expression plasmids were transformed into *E. coli* BL21 DH5*α*, spread on LB agar plates containing 50 μg·mL^-1^ kanamycin or 100 μg·mL^-1^ ampicillin and incubated (37°C, ON). Appearing single colonies were inoculated into 3 mL LB medium with appropriate antibiotics and cultivated (37°C, ON, 150 rpm) followed by plasmid purification. The gene inserts in purified plasmids were verified by Sanger sequencing (Eurofins Genomics, Germany) and transformed into *E. coli* BL21 (DE3) (New England Biolabs). In an attempt to improve transglycosylation, the mutants Fp231_H174F and Fp231_W225H were designed based on mutant transfer (Teze et al. 2021) and introduced into the Fp231 gene as described (Liu and Naismith 2008) using the primers listed in **Supplementary Table I**.

### Production and purification of recombinant enzymes

*E. coli* BL21 (DE3) transformants with pET15b expression plasmids were inoculated in 50 mL ZYP autoinduction medium (Studier 2005) supplemented with 100 μg·mL^-1^ ampicillin, incubated (20°C, 72 h, 150 rpm), centrifuged (20,000 x *g*, 10 min, 4°C) and the pellets stored at −20°C. Thawed cells were resuspended in 10 mL 10 mM imidazole, 50 mM HEPES pH 7.5, 300 mM NaCl (IMAC A buffer) and lysed using a pressure cell homogenizer (Stansted). Cell debris was removed by centrifugation (20,000 x *g*, 20 min) and the supernatants were agitated with 300 μL Ni^2+^-NTA resin using a magnetic stirring bar (4°C, 15 min). The Ni^2+^-NTA resin was transferred to empty Econo-Pac® columns (Bio-Rad), washed with 15 mL IMAC A buffer containing 25 mM imidazole followed by elution with 1.6 mL IMAC A buffer containing 250 mM imidazole.

*E. coli* BL21 (DE3) transformed with pET9a for expression of wild type Fp231, Fp231_H174F or Fp231_W225H was cultivated in LB medium containing 100 μg·mL^-1^ kanamycin (37°C, ON, 150 rpm). This overnight culture was diluted 1:200 in fresh medium (6 x 1 L) and cultivated (37°C, 150 rpm) until an OD_600_ = 0.6–0.8, and induced by addition of isopropyl β-D-1-thiogalactopyranoside (IPTG, 0.1 mM final concentration, 20°C, 18 h, 150 rpm). Cells were harvested by centrifugation (15,000 x *g*, 40 min, 4°C) and the supernatant was filtered (0.2 μm, Millipore) before concentration on a tangential flow filtration system equipped with a 30 kDa cut-off filter membrane (Sartorius). The ~200 mL retentate was equilibrated to 20 mM imidazole, 20 mM Tris-HCl, pH 7.5, 300 mM NaCl (IMAC B buffer) and loaded onto a 5 mL HiTrap IMAC column (GE Healthcare) using an Äkta Start Chromatography system. The column was washed with 5 column volumes IMAC B buffer containing 100 mM imidazole and Fp231 was eluted by 5 column volumes IMAC B buffer containing 250 mM imidazole. Fp231 was further purified by gel filtration (HiPrep™ 16/60 Sephacryl™ S-200 HR; GE Healthcare, Uppsala, Sweden) in 50 mM sodium citrate-phosphate pH 6.0, 150 mM NaCl. Protein purity was assessed by SDS-PAGE and analytical size exclusion chromatography (ENrich SEC 650 column, BioRad) using a Gel Filtration Standard for calibration (Bio-Rad, #1511901). Purified enzyme was concentrated (30 kDa cutoff Amicon Stirred Cell, Millipore or VivaSpin4 columns, Sartorius). Protein concentration was determined spectrophotometrically at 280 nm (μCuvette, Eppendorf) using a theoretical extinction coefficient for Fp231 of 110935 M^-1^·cm^-1^ (ProtParam: https://web.expasy.org/protparam/)

### Activity assay and substrate specificity screening

*α*-l-fucosidase activity was measured towards 1 mM 2-chloro-4-nitrophenyl-*α*-l-fucopyranoside (CNP-Fuc) by the increase in absorbance at 410 nm corrected by a control reaction mixture with heat-inactivated enzyme and using an extinction coefficient for CNP determined to 11283 M^-1^·cm^-1^ in citrate-phosphate pH 6.0. For substrate screening of enzymes, purified recombinant Fp231, Fp239, Fp240, Fp251 or Fp284 (>0.2 μM) was mixed with 2 mM oligosaccharide in 10 μL 20 mM sodium citrate-phosphate pH 6.2, then incubated for 6–24 h at 37°C and heat-inactivated (98°C, 5 min). Samples were diluted 1:10 in H_2_O and analyzed on a Dionex 3000 HPLC system equipped with a PA-1 analytical column. The elution was performed with 75 mM NaOH in sodium acetate 0.1 M at 250 μL·min^-1^ for 20 min.

### Biochemical characterization of Fp231

pH and temperature optima and suitable buffer conditions were determined using 1 mM CNP-Fuc as substrate and 5 nM Fp231. Activity was assayed in 50 mM citrate/100 mM phosphate (pH 4.0 – 8.0) or in 50 mM MES, Bis-Tris, MOPS and Tris-HCl in the range pH 6.0 – 7.0 for 20 min in the range 5 – 40°C using a thermocycler. Effect of 0 – 2 M NaCl on activity was determined at the optimal pH and temperature. The effect of metal ions on activity was tested by assaying Fp231 in the presence of 1 mM or 5 mM CaCl_2_, ZnCl_2_, CoCl_2_, NiCl_2_, MnCl_2_, MgCl_2_, LiCl, KCl or EDTA in Bis-Tris pH 6.5.

Enzyme reactions (100 μL) for fluorophore-assisted carbohydrate electrophoresis (Starr et al. 1996) containing 5 μM Fp231 and 10 μg oligosaccharide or 1% fucoidan were incubated (R.T., ON) and heat-inactivated (99°C, 10 min). Samples were dried in a vacuum concentrator centrifuge (45 min, 60°C) and dissolved in 2 μL 8-aminonaphthalene-1,3,6-trisulfonic acid fluorophore (0.2 M in 3:17 acetic acid:H_2_O v/v) and 5 μL NaBH3CN (1 M in tetrahydrofuran) followed by incubation (37°C, 18 h). The fluorescently labeled reaction products were separated in acrylamide gels (27 or 35 % resolving/4 % stacking) for 1 h at 100 V, then 1 h at 300 V, and visualized under UV light.

Initial rates of hydrolysis of 0.1 – 4 mM CNP-Fuc were determined using 2 nM Fp231 and the formation of CNP was monitored every 30 s over 30 min. Initial rates of hydrolysis of 0.1 – 4 mM Fuc(*α*1,4)GlcNAc were determined from the fucose released by 2 nM Fp231 sampled at 15, 30, 45 and 60 min, analyzed using by HPAEC-PAD as described (McGregor et al. 2017). Kinetic constants were calculated from Michaelis-Menten plots with non-linear regression using an R script as described (Huitema and Horsman 2018).

Transglycosylation activity of wildtype Fp231 and mutants was tested by incubating 20 mM CNP-Fuc and 200 mM GlcNAc with 5 – 500 nM enzyme in citrate-phosphate at 25 °C for up to 24 h. The formation of transglycosylation products was analyzed by FACE as described above.

### Protein crystallization trials

Crystallization was attempted by the sitting drop vapor diffusion method of 20–80 mg·mL^-1^ purified Fp231 with a selection of commercial screens (**Supplementary Table II**). A Phoenix pipetting robot (Art Robbins) was used to dispense 96-well plates with 80 μL mother liquor in reservoirs and 300 nL droplets containing mother liquor and protein in the ratio of 1:1 or 1:2 (vol/vol). Plates were covered with adhesive film and kept at 16°C and inspected regularly during a 12 month period.

## Supporting information

Supplemental Tables SI-SII and Figures S1-S6

## Acknowledgements

Prof. Jan-Hendrik Hehemann is acknowledged for valuable discussions and for providing additional funding through the Emmy-Noether Program of the German Research Foundation [HE 7217/1-1]. Morten Jensen is gratefully acknowledged for assistance with cloning and screening of enzymes. We thank Karina Jansen for technical support.

## Supplementary data

Figures S1 – S6 and Tables SI – SII are included in this manuscript as a single pdf file.

## Abbreviations

CAZy: Carbohydrate-active enzyme
CNP-Fuc: 2-chloro-4-nitrophenyl-*α*-l-fucopyranoside
CSDB: Carbohydrate structure database
FACE: Fluorophore-assisted carbohydrate electrophoresis
FucGlcNAc: Fucosyl-*N*-acetylglucosamine
GH29: Glycoside hydrolase family 29
HMO: Human milk oligosaccharide
HMW: High molecular weight
HPAEC-PAD: High pressure anion exchange chromatography coupled with pulsed amperometric detection
IMAC: Immobilized metal affinity chromatography
LMW: Low molecular weight
SEC: Size exclusion chromatography

## Data availability

The data underlying this article are available in the GenBank Nucleotide Database at https://www.ncbi.nlm.nih.gov/genbank/, and can be accessed under accession numbers MW623630 (Fp231), MW623631 (Fp239), MW623632 (Fp240), MW623633 (Fp251) and MW623634 (Fp284).

