## Supplemental Tables SI-SII and Figures S1-S6 for "Two marine GH29 *α*-L-fucosidases from an uncultured *Paraglaciecola* sp. specifically hydrolyze fucosyl-*N*-acetylglucosamine regioisomers"

### Supplementary Table I:

Nucleotide primers used in the present study. Uracil-containing overhangs for USER-fusion are underscored and codon triplets changed for mutants are in bold.

| **Primer name** | **5'-->3'** |
|---|---|
| *Primers for pET15b cloning* | |
| Fp231_pET15b_F | ACG GAT CUG AGC TTG CCG CCC AC |
| Fp231_pET15b_R | AGC CGG AUT TAC TGG CTT ATC TCT AAT GC |
| Fp239_pET15b_F | ACG GAT CUC AAG CAC ACT CTG AAA AAA C |
| Fp239_pET15b_R | AGC CGG AUC TAA GGG AGA TAT TCT ATA GTT AAC |
| Fp240_pET15b_F | ACG GAT CUG CCG ATA AAC CCT ACG AC |
| Fp240_pET15b_R | AGC CGG AUT TAA AAA TGA TTT CCG TCT AAA TC |
| Fp251_pET15b_F | ACG GAT CUG AGA CTG AGC ATA GAC TTA AAC |
| Fp251_pET15b_R | AGC CGG AUT TAC TTT AAT AAC ACT TCA ATG G |
| Fp284_pET15b_F | ACG GAT CUG CAA GTG ATT ATA CAA GCC TTA C |
| Fp284_pET15b_R | AGC CGG AUC TAC TGC GCT TCT TTA ACC |
| *Primers for pET9a cloning* | |
| Fp231_Nat.SP_Forw. | AGG CTT AAU ATG CAT CAG CAA CGA G |
| Fp231_Nat.SP_Rev. | ACT TCC ACU CTG GCT TAT CTC TAA TGC |
| Fp231_K67_Forw. | ATA TGG CUA AAA GCT TGA CCA AAG AG |
| Fp231_K417_Rev. | ACT TCC ACU TTT AGA GTC ATA GAT TGC ATC |
| Fp231_E48_Forw. | AGG CTT AAU GAG AAA AAA CAA GTA TAT GGC |
| *Primers for mutagenesis* | |
| Fp231_H174F_Forw. | GCT CCA AG**T TC**C ATG AAG GTT TTG CCA TGT TTA AGT CTG AAG |
| Fp231_H174F_Rev. | TTC ATG **GAA** CTT GGA GCC CAT GAC GAT GTA TTT CAT GCC T |
| Fp231_W225H_Forw. | AAC TCC CTT GAT **CAT** CGC GAT GGT GGT GAT GGT GG |
| Fp231_W225H_Rev. | CG**A TG**A TCA AGG GAG TTA GAA TAA TAA ACA CCA AAG TCG AGG C |
| *Primers for vector amplification* | |
| pET15b_vec_Forw. | ATC CGG CUG CTA ACA AAG |
| pET15b_vec_Rev. | AGA TCC CUG ATG ATG ATG ATG ATG GCT |
| pET9a_vec_Forw. | ATT AAG CCU CAG CAT ATG TAT ATC |
| pET9a_vec_Rev. | AGT GGA AGU CCG CAC CAC CAC CAC |

**Supplementary Table II:** 96-well format protein crystallization screens used in this study.

| **Screen** | **Supplier** |
|---|---|
| MCSG-1 | Anatrace |
| MCSG-2 | Anatrace |
| SaltRx 1 | Hampton Research |
| JBScreen Basic HTS | Jena Biosciences |
| JBScreen Pact ++ HTS | Jena Biosciences |
| JBScreen Clasic HTS I | Jena Biosciences |
| JBScreen PEG/Salt HTS | Jena Biosciences |
| JBS JCSG++ HTS | Jena Biosciences |
| The PGA Screen™ (MD1-51) | Molecular dimensions |
| SG1™ Screen (MD1-89) | Molecular dimensions |
| Clear Strategy™ (MD1-31) | Molecular dimensions |
| Morpheus® (MD1-47) | Molecular dimensions |
| MIDAS*plus*™ (MD1-107) | Molecular dimensions |

**Figure S1:** Protein domain structure of GH29 fucosidases in the present study. Protein domain features were predicted using the hmmscan function at EMBL-EBI https://www.ebi.ac.uk/ and SignalP 5.0 (Almagro Armenteros et al. 2019). The total length of each enzyme is indicated in amino acids (AA). SPI: Secretory signal peptide cleaved by Signal Peptidase I, SPII: Lipoprotein signal peptide cleaved by Signal Peptidase II

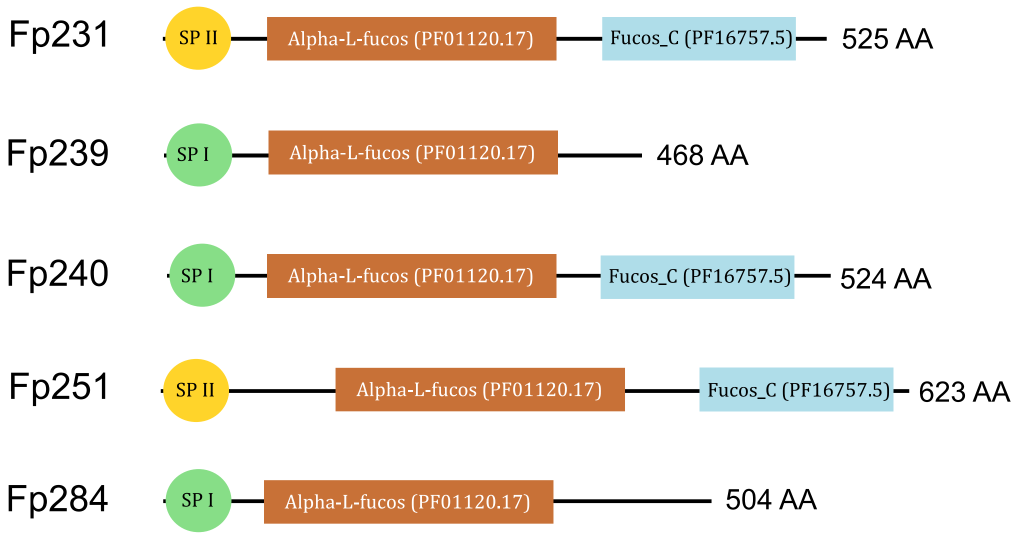

**Figure S2:** SDS-PAGE of recombinant GH29 fucosidases after IMAC purification. Theoretical molar masses: Fp239 (52.4 kDa), Fp240 (59.4 kDa), Fp251 (70.3 kDa), and Fp284 (56.8 kDa).

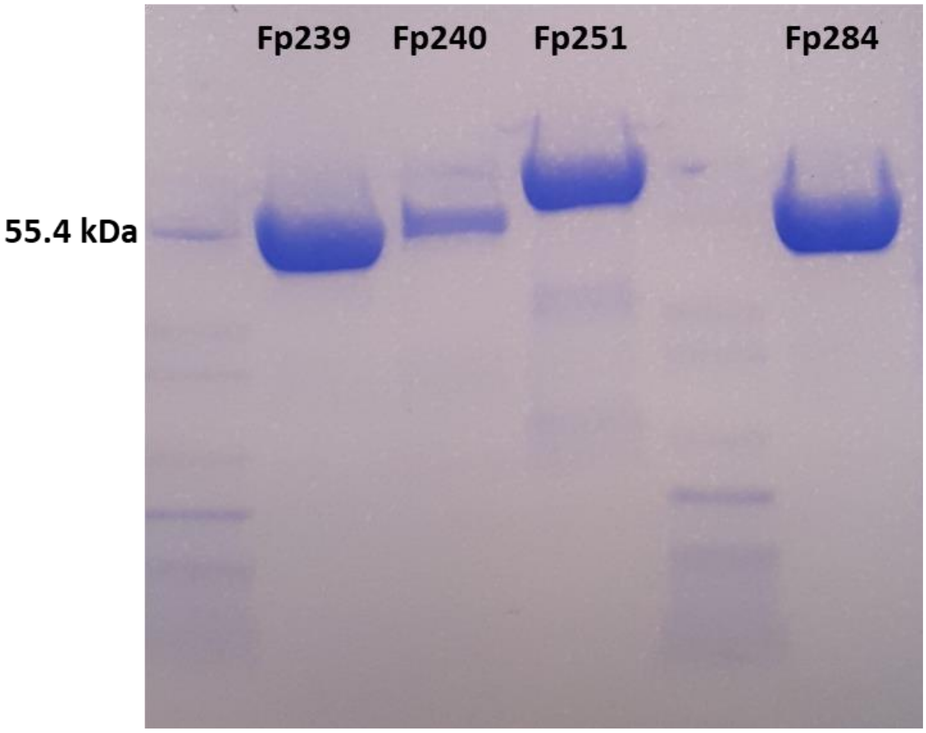

**Figure S3:** Activity of Fp231 in the presence of NaCl measured towards 1 mM CNP-Fuc.

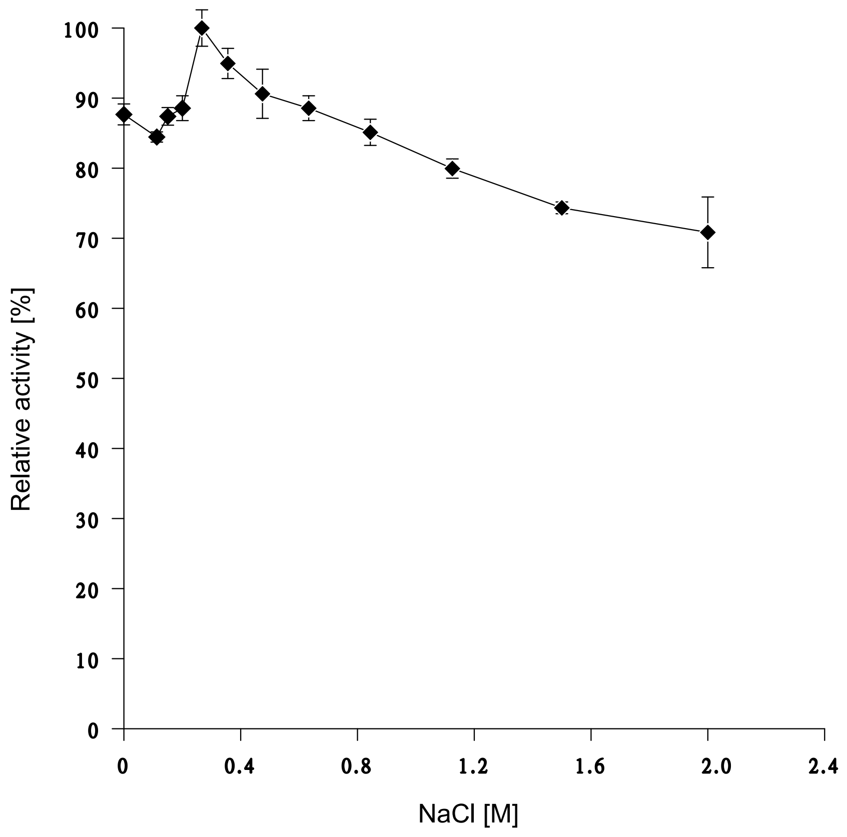

**Figure S4:** Michaelis-Menten plots of hydrolysis of CNP-Fuc and Fuc($\alpha$1,4)GlcNAc by Fp231

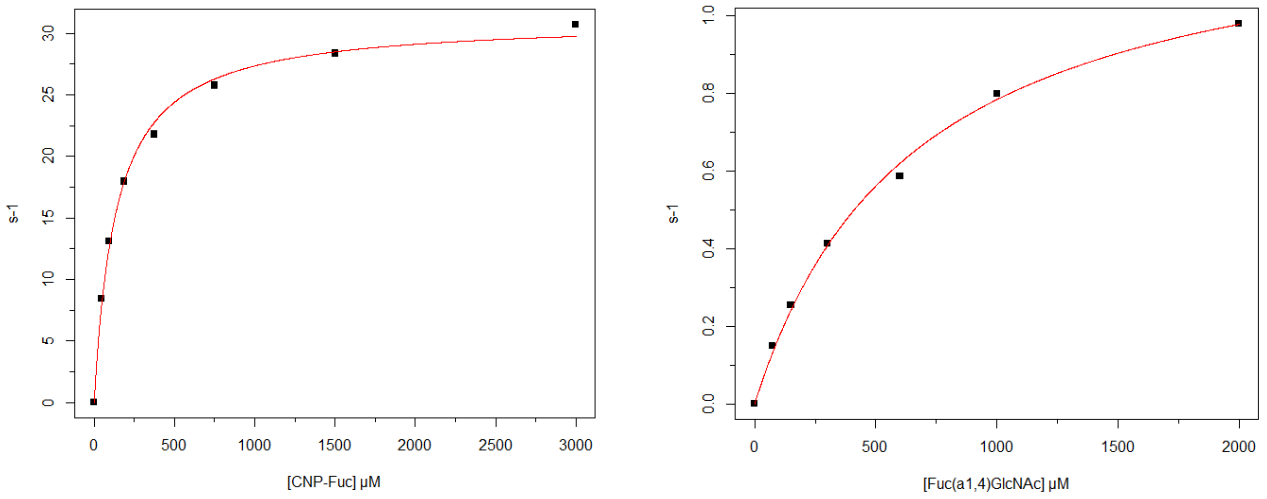

**Figure S5:** FACE gels showing activity of Fp231 on oligosaccharides (supplementing Figure 3B). Oligosaccharide substrates (**A** and **B**), or algal fucoidans from *Fucus vesiculosus, Fucus serratus* and *Ascophyllum nodosum* (**B**), were incubated with (+) and without (–) Fp231 (5 µM) at room temperature for 24 h and the reaction products were fluorescently labeled and separated in acrylamide gels. Enzyme activity was evaluated as a change in mobility of the fluorescent products. Monosaccharide standards (Fuc and GlcNAc) are included for comparison.

**A)**

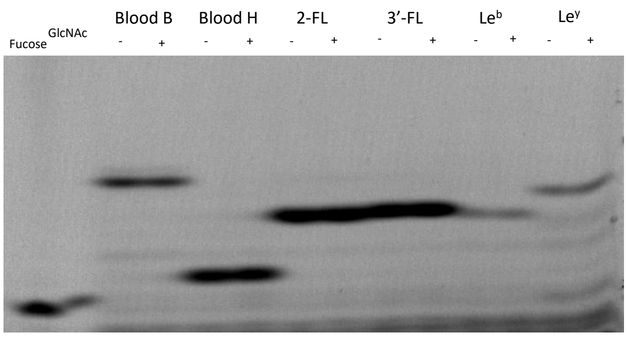


**B)**

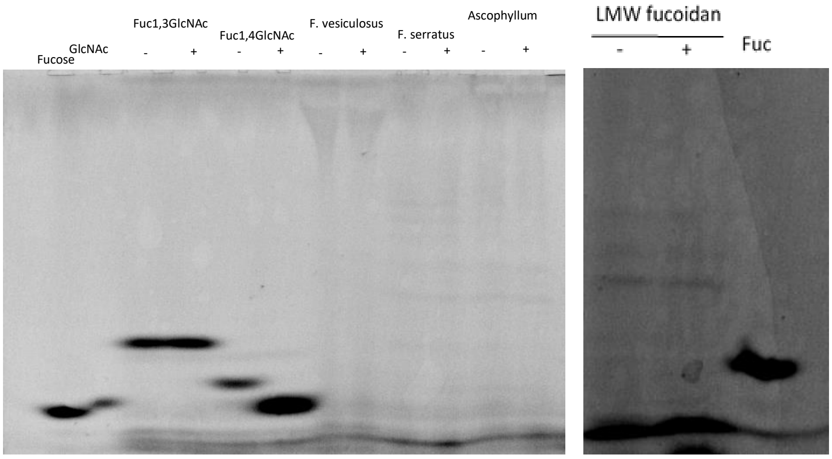

**Figure S6:** Overlay of oligosaccharide structures. **A)**: Fuc($\alpha$1,4)Gal (green sticks) and Fuc($\alpha$1,4)Fuc (cyan sticks); **B)**: Fuc($\alpha$1,4)GlcNAc (pink sticks), Fuc($\alpha$1,4)Gal (green sticks) and Fuc($\alpha$1,4)Fuc (cyan sticks)

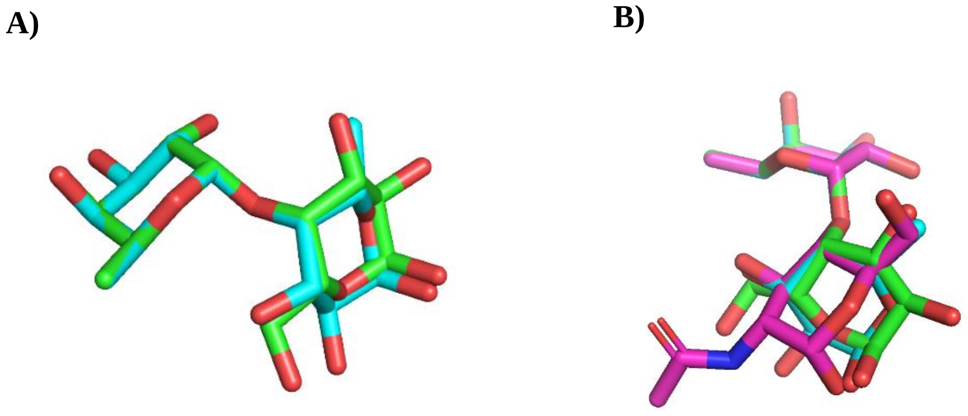
